# ERK3-MK5 signaling regulates myogenic differentiation and muscle regeneration by promoting FoxO3 degradation

**DOI:** 10.1101/2021.04.29.441967

**Authors:** Mathilde Soulez, Pierre-Luc Tanguay, Florence Dô, Colin Crist, Junio Dort, Alexey Kotlyarov, Matthias Gaestel, Nicolas A. Dumont, Sylvain Meloche

## Abstract

The physiological functions and downstream effectors of the atypical mitogen-activated protein kinase ERK3 remain to be characterized. We recently reported that mice expressing catalytically-inactive ERK3 (*Mapk6*^*KD/KD*^) exhibit a reduced post-natal growth rate as compared to control mice. Here, we show that genetic inactivation of ERK3 impairs post-natal skeletal muscle growth and adult muscle regeneration after injury. Loss of MK5 phenocopies the muscle phenotypes of *Mapk6*^*KD/KD*^ mice. At the cellular level, genetic or pharmacological inactivation of ERK3 or MK5 induces precocious differentiation of C2C12 or primary myoblasts, concomitant with MyoD activation. Reciprocally, ectopic expression of activated MK5 inhibits myogenic differentiation. Mechanistically, we show that MK5 directly phosphorylates FoxO3, promoting its degradation and reducing its association with MyoD. Depletion of FoxO3 rescues in part the premature differentiation of C2C12 myoblasts observed upon inactivation of ERK3 or MK5. Our findings reveal that ERK3 and its substrate MK5 act in a linear signaling pathway to control post-natal myogenic differentiation.

## INTRODUCTION

ERK3 (Extracellular Signal-Regulated Kinase 3) is an atypical member of the MAPK (mitogen-activated protein kinase) family that is ubiquitously expressed in mammalian tissues (1). Much remains to be learned about the physiological functions of ERK3. The expression of ERK3 is temporally regulated during embryogenesis, increasing at the time of early organogenesis and then declining gradually toward birth (2). Early studies showed that ERK3 expression is upregulated during in vitro differentiation of model cell lines into neurons and muscle cells (3, 4). These observations point to a possible role of ERK3 signaling in cellular differentiation. Genetic invalidation studies have revealed that mice lacking ERK3 expression or activity are born at normal mendelian frequency and survive to adulthood without any apparent health problem (5, 6). However, ERK3 kinase activity is required for optimal post-natal growth of newborn mice (6).

Little is known about the repertoire of ERK3 substrates, which has limited our understanding of its cellular functions. Unlike classical MAPKs like ERK1/2 that phosphorylate a large spectrum of substrates, ERK3 has a more restricted substrate specificity. Its only validated substrate is the protein kinase MAPK-activated protein kinase 5 (MK5) (7, 8). ERK3 binds to and phosphorylates MK5 on the activation loop Thr182, leading to its enzymatic activation and to cytoplasmic relocalization of the complex. The relative contribution of MK5 to the physiological functions of ERK3 remains to be established. In hippocampal neurons, ERK3 forms a complex with MK5 and the cytoskeletal protein septin 7, and co-expression of these proteins increases neuron branching and spine number, suggesting a possible role of ERK3-MK5 signaling in neuronal morphogenesis (9). A recent study reported that β-adrenergic-stimulated protein kinase A directly phosphorylates MK5, leading to the formation of a stable ERK3-MK5 complex which promotes lipolysis by inducing the expression of the major lipase ATGL (10). The exact role of MK5 in this ERK3-dependent lipolytic signaling pathway is not fully understood.

Skeletal muscle is the most abundant tissue of the body, accounting for ∼40% of body weight (11). The development of skeletal muscle, or myogenesis, occurs in multiple phases of growth that mobilize stem and progenitor cells to establish the body skeletal muscles (12–14). In the mouse, skeletal muscle is established from E8.5 to E18.5, with further maturation during the first 3 weeks of postnatal life. In the perinatal phase, muscle resident Pax7^+^ progenitors proliferate extensively to provide new myonuclei to secondary myofibers, after which proliferation gradually ceases and myofibrillar protein synthesis reaches a peak. The signaling events controlling post-natal skeletal muscle growth remain to be fully defined. Once the muscle has matured, cycling Pax7^+^ progenitors enter quiescence and reside along host myofibers as adult satellite cells. Upon muscle injury, satellite cells are rapidly activated to re-enter the cell cycle, generating proliferating myoblasts that expand through several rounds of division before entering the myogenic differentiation program (15, 16). Terminally differentiated myocytes then fuse to each other or to damaged fibers to form new myofibers and re-establish homeostasis. Developmental myogenesis and muscle regeneration are instructed by a spectrum of extrinsic factors and signaling pathways that converge on a hierarchical transcriptional regulatory network comprising the core myogenic regulatory factors Myf5, MyoD, Mrf4 and myogenin (12, 13, 17–19).

In this study, we tested the hypothesis that ERK3 is involved in the regulation of skeletal muscle development. We show that mice expressing a kinase-dead allele of ERK3 have defective post-natal myogenesis and muscle regeneration after injury. Genetic invalidation of MK5 essentially phenocopies the muscle phenotypes of ERK3-inactive mice. At the cellular level, inactivation of ERK3 or MK5 induces premature differentiation of proliferating myoblasts into myotubes. We further show that MK5 phosphorylates FoxO3, which promotes its degradation and decreases its association with MyoD. Our findings unveil a new function of ERK3-MK5 signaling in regulating post-natal myogenic differentiation.

## RESULTS

### Defective post-natal skeletal muscle growth in mice expressing catalytically-inactive ERK3

We have shown previously that mice expressing a kinase-dead (KD) allele of ERK3 (*Mapk6*^*KD/KD*^) have a reduced post-natal growth rate as compared to wild type (WT) control mice (6). This growth phenotype affected both female and male mice. No difference in body weight was observed between the two genotypes at birth (6). Given the close correlation between the increase in body mass and muscle mass during the first weeks of post-natal life (20), we measured the cross-sectional area (CSA) of individual myofibers from the entire tibialis anterior (TA) muscle of littermate WT and *Mapk6*^*KD/KD*^ mice at post-natal day (P) 15. The average size of TA myofibers was significantly smaller in *Mapk6*^*KD/KD*^ mice with a shift in myofiber CSA toward smaller fibers (Fig. 1 A and B). There was no change in the total number of myofibers per TA muscle between WT and *Mapk6*^*KD/KD*^ mice (Fig. 1 C and D), suggesting normal specification of skeletal myoblasts. We conclude that ERK3 kinase activity is required for post-natal skeletal muscle growth.

**Figure 1.**
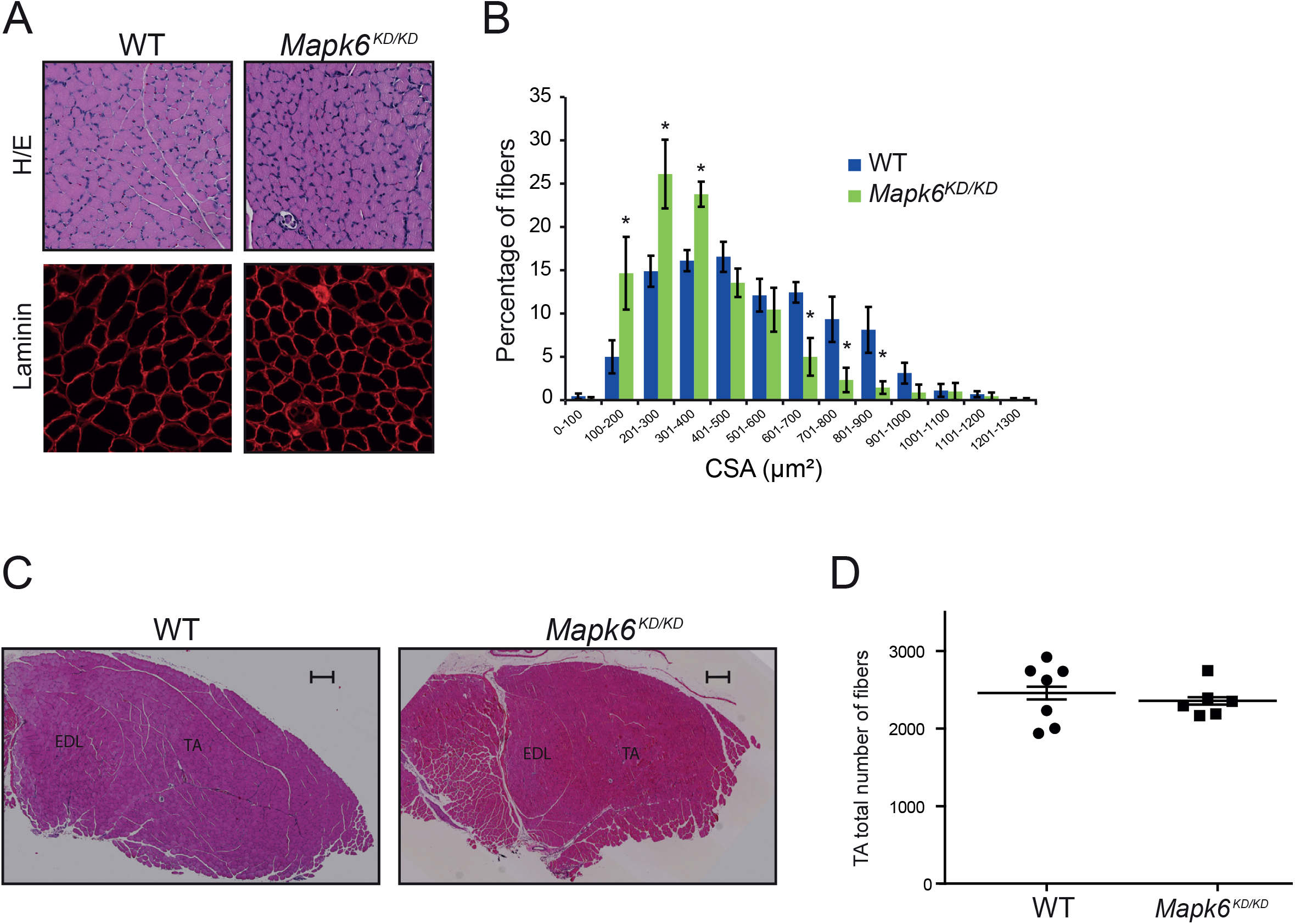
ERK3 kinase activity regulates post-natal skeletal muscle growth in mice. (A) Sections of TA muscle from WT and *Mapk6*^*KD/KD*^ mice at P15 analyzed by H&E staining and laminin immunostaining. Scale bar, 100 µm. (B) Distribution of myofiber CSA in TA muscle of WT and *Mapk6*^*KD/KD*^ mice at P15. Values are mean ± SEM (n = 6). \**p* < 0.05. (C) H&E staining of WT and *Mapk6*^*KD/KD*^ mice TA muscle at 10 weeks. Scale bar, 300 µm. (D) Analysis of total myofiber content in TA muscle of WT and *Mapk6*^*KD/KD*^ mice at 10 weeks. Values are mean ± SEM (n Λ6).

### Genetic inactivation of ERK3 impairs skeletal muscle regeneration

We next investigated the function of ERK3 in adult myogenesis using a model of skeletal muscle regeneration after cardiotoxin (CTX)-induced injury (21). For these experiments, CTX was injected into the TA muscle of adult mice, and the injured muscles were analyzed at different times during the regeneration process (Supplementary Fig. S1). We found that ERK3 mRNA and protein levels are transiently up-regulated during muscle regeneration, reaching a peak at ∼ 3 days and declining thereafter (Fig. 2 A and B), coincident with the proliferation and expansion of myoblasts (16, 22). To address the role of ERK3, we compared the regenerating capacity of WT and *Mapk6*^*KD/KD*^ mice after CTX injury. Genetic inactivation of ERK3 impaired muscle regeneration as revealed by a significant decrease in myofiber CSA 15 days post-injury (Fig. 2 C and D). Since muscle regeneration is absolutely dependent on adult satellite cells (23–25), we quantified the number of Pax7^+^ satellite cells in the TA muscle. No difference in the number of Pax7^+^ cells was observed between WT and *Mapk6*^*KD/KD*^ mice (Fig. 2 E and F), consistent with the idea that specification of Pax7^+^ progenitors during fetal myogenesis is normal in mice expressing ERK3 KD. Thus, ERK3 kinase activity is required for normal muscle regeneration after injury.

**Figure 2.**
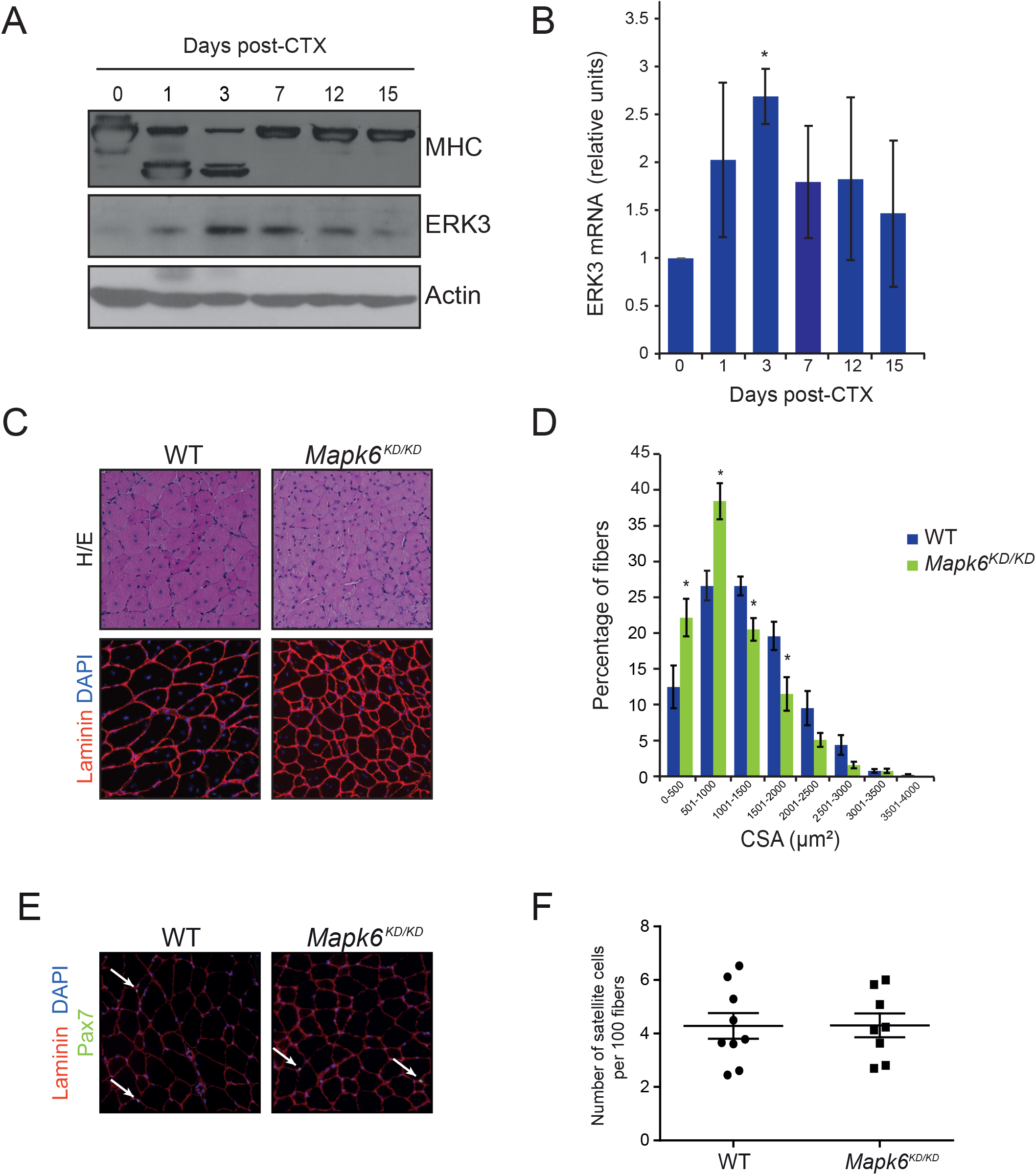
ERK3 kinase activity regulates injury-induced skeletal muscle regeneration. (A) Expression of ERK3 during skeletal muscle regeneration. TA muscle of 10 week-old mice was injured by CTX injection and harvested at the indicated times. Expression of ERK3 and MHC was analyzed by immunoblotting in total lysates. (B) Expression of ERK3 mRNA during muscle regeneration was measured by qPCR. Values are mean ± SEM (n = 3). \*\**p* < 0.01. (C) Sections of TA muscle from WT and *Mapk6*^*KD/KD*^ mice analyzed by H&E staining, laminin immunostaining and DAPI 15 days post-CTX injection. Scale bar, 100 µm. (D) Distribution of myofiber CSA in TA muscle of WT and *Mapk6*^*KD/KD*^ mice 15 days post-injury. Values are mean ± SEM (n Λ7). \**p* < 0.05. (E) Immunofluorescence staining of laminin (red) and Pax7 (green) in TA muscle of WT and *Mapk6*^*KD/KD*^ mice at 10 weeks. (F) Analysis of Pax7^+^ satellite cells in TA muscle of WT and *Mapk6*^*KD/KD*^ mice at 10 weeks. Values are mean ± SEM (n Λ8).

### Loss of MK5 phenocopies the muscle defects observed in mice expressing catalytically-inactive ERK3

We then asked whether MK5 contributes to the regulation of post-natal growth and skeletal muscle development by ERK3 in mice. The growth rate of *Mapkakpk5^−/−^* homozygous mice was clearly reduced as compared to littermate control or heterozygous mice during the first two weeks of post-natal life (Fig. 3A). Analysis of TA myofiber size at P15 showed that myofiber CSA is significantly shifted toward smaller fibers in *Mapkakpk5^−/−^* mice (Fig. 3 B and C). In the CTX model of muscle regeneration, loss of MK5 impaired the regeneration process as characterized by smaller myofibers 15 days after injury (Fig. 3D and E). Thus, loss of MK5 phenocopies the muscle defect phenotypes of ERK3 KD-expressing mice, suggesting that ERK3 and MK5 are parts of the same signaling pathway to regulate post-natal skeletal muscle development.

**Figure 3.**
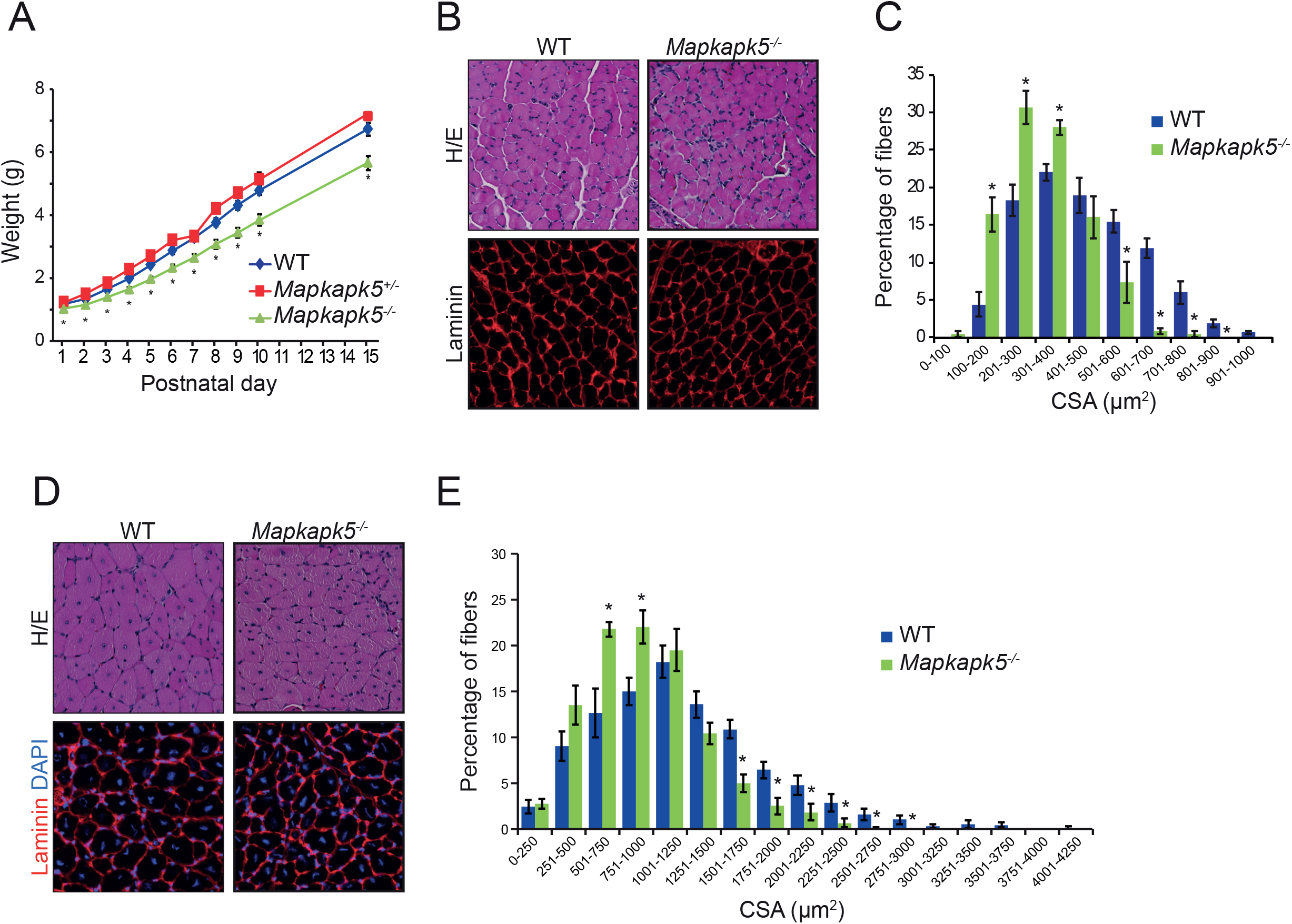
Loss of MK5 phenocopies the muscle defects of mice expressing kinase-dead ERK3. (A) Growth curves of WT, *Mapkapk5*^*+/−*^ and *Mapkapk5^−/−^* mice over 15 post-natal days. Values are mean ± SEM (n Λ15). \**p* < 0.05. (B) Sections of TA muscle from WT and *Mapkapk5^−/−^* mice at P15 analyzed by H&E staining and laminin immunostaining. Scale bar, 100 µm. (C) Distribution of myofiber CSA in TA muscle of WT and *Mapkapk5^−/−^* mice at P15. Values are mean ± SEM (n Λ3). \**p* < 0.05. (D) Sections of TA muscle from WT and *Mapkapk5^−/−^* mice analyzed by H&E staining, laminin immunostaining and DAPI 15 days after CTX injection. Scale bar, 100 µm. (E) Distribution of myofiber CSA in TA muscle of WT and *Mapkapk5^−/−^* mice 15 days post-injury. Values are mean ± SEM (n Λ6). \**p* < 0.05.

### Inactivation of ERK3 or MK5 leads to precocious myogenic differentiation of skeletal myoblasts

To understand the cellular mechanisms underlying the skeletal muscle growth defect observed upon inactivation of ERK3-MK5 signaling, we depleted ERK3 from mouse C2C12 myoblasts (26) by CRISPR/Cas9 gene editing. Using two distinct sgRNAs, we generated populations of C2C12 myoblasts with >90% depletion of ERK3 (Fig. 4A and Supplementary Fig. S2A). Loss of ERK3 had no significant effect on the rate of proliferation of C2C12 myoblasts (Supplementary Fig. S2B). We then analyzed the ability of C2C12 cell populations to differentiate into myotubes in vitro. Depletion of ERK3 induced premature differentiation of C2C12 myoblasts as revealed by increased expression of the terminal differentiation marker myosin heavy chain (MHC) after 1 to 3 days in differentiation medium (DM) and by up-regulation of myogenin levels in growth medium (GM) and at DM1 (Fig. 4A and Supplementary Fig. S2A). Likewise, immunofluorescence analysis showed that ERK3-depleted C2C12 myoblasts form larger myotubes and display a higher fusion index as compared to parental cells (Fig. 4 B and C and Supplementary Fig. S2C). No further increase in MHC expression or fusion index was observed when both ERK3 and MK5 were depleted from C2C12 cells, suggesting that the two kinases act in the same pathway (Supplementary Fig. S2 A and C). Essentially similar results were obtained when ERK3 expression was silenced by a lentiviral shRNA (Supplementary Fig. S2D). To confirm these findings, we isolated satellite cell-derived primary myoblasts from WT and *Mapk6*^*KD/KD*^ mice and analyzed their ability to differentiate in cell culture. Genetic inactivation of ERK3 markedly reduced the proliferation rate of primary myoblasts (Fig. 4D), and accelerated their differentiation into myotubes characterized by the formation of longer and thicker myotubes and higher fusion index (Fig. 4 E and F and Supplementary Fig. S2E). These results indicate that ERK3 kinase activity is required to prevent premature myogenic differentiation of myoblast into myotubes.

**Figure 4.**
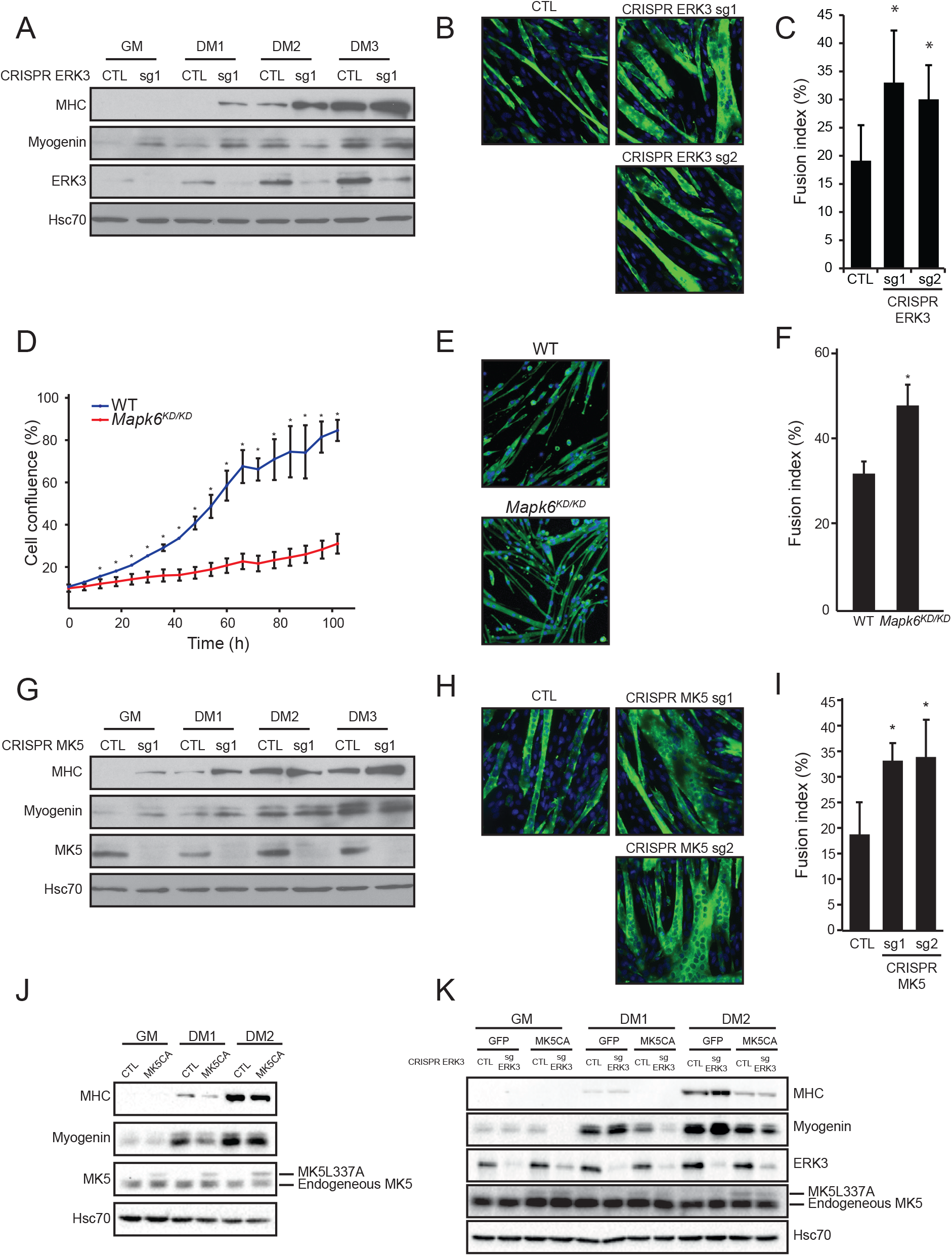
ERK3 and MK5 lie on a linear signaling pathway to control differentiation of skeletal myoblasts. (A-C) Analysis of C2C12 myoblasts depleted of ERK3 by CRISPR/Cas9 gene editing. C2C12 cells were transfected with empty vector (CTL) or PX459 vector expressing sgRNA sequences (sg) targeting *Mapk6* gene. Stable populations were generated by puromycin selection and used thereafter. The cells were grown in GM medium and switched to DM medium to initiate myogenic differentiation. (A) Expression of MHC, myogenin and ERK3 was analyzed by immunoblotting at the indicated times. (B) Immunofluorescence staining of MHC expression at DM3. (C) Quantification of fusion index of differentiating C2C12 myoblasts at DM3. Values are mean ± SD (n Λ4). \**p* < 0.05. (D-F) Analysis of satellite cell-derived primary myoblasts isolated from WT and *Mapk6*^*KD/KD*^ mice. Myogenic differentiation was induced by replacing GM with DMEM containing 5% horse serum. (D) Cell proliferation was monitored using an automated IncuCyte live-cell imaging system. Values are mean ± SD (n=3). \**p*<0.05. (E) Immunofluorescence staining of MHC expression 4 days after induction of differentiation. (F) Quantification of fusion index of differentiating primary myoblasts 4 days after differentiation induction. Values are mean ± SEM (n = 3). \**p* < 0.05. (G-I) Analysis of C2C12 myoblasts depleted of MK5 by CRISPR/Cas9 gene editing. (G) Expression of MHC, myogenin and MK5 was analyzed by immunoblotting at the indicated times. (H) Immunofluorescence staining of MHC expression at DM3. (I) Quantification of fusion index of differentiating C2C12 myoblasts at DM3. Values are mean ± SD (n Λ3). *p < 0.05. (J) Effect of constitutively active (CA) MK5 L337A on C2C12 myoblast differentiation. C2C12 cells were transfected with MK5 L337A. Stable populations were generated by selection in 500 µg/ml G418 for 10 days. The cells were grown in GM and switched to DM to initiate myogenic differentiation. Expression of MHC, myogenin and MK5 was analyzed by immunoblotting. (K) Activated MK5 L337A was expressed in C2C12 myoblasts depleted of ERK3 by CRISPR/Cas9 editing. Expression of MHC, myogenin, ERK3 and MK5 was analyzed by immunoblotting.

We used CRISPR/Cas9 editing to generate populations of C2C12 myoblasts with an almost complete depletion of MK5 (Fig. 4G and Supplementary Fig. S2A). Loss of MK5 did not affect the proliferation rate of C2C12 cells (Supplementary Fig. S2B). Similar to ERK3, depletion of MK5 induced premature differentiation of C2C12 myoblasts into myotubes (Fig. 4 G-I and Supplementary Fig. S2 A and C). Comparable results were obtained when depleting MK5 by RNA interference (Supplementary Fig. S2F) or by inhibition of MK5 kinase activity with the chemical inhibitor PF-3644022 (27) (Supplementary Fig. S2G-I). Conversely, overexpression of a constitutively active (CA) mutant of MK5, MK5 L337A (28), in C2C12 myoblasts inhibited myogenic differentiation (Fig. 4J). To further address the functional relationship of ERK3 and MK5 in the myogenic differentiation process, we ectopically expressed MK5 L337G in ERK3-depleted C2C12 cells. Despite the relatively weak level of MK5 L337A expression (compared to endogenous MK5), we found that activation of MK5 prevented the premature differentiation of C2C12 myoblasts induced by the loss of ERK3 (Fig. 4K). These results provide compelling evidence that ERK3 and MK5 act in the same pathway to restrict myogenic differentiation.

### ERK3-MK5 signaling regulates MyoD activity in myoblasts

To gain insight into the molecular mechanism by which ERK3 and MK5 control myogenic differentiation, we compared the gene expression profiles of parental C2C12 myoblasts with that of C2C12 cells depleted of ERK3 or MK5 by CRISPR/Cas9 editing in GM and at DM2. Principal component analysis (PCA) showed that control, ERK3-depleted and MK5-depleted C2C12 cells are grouped into one cluster at GM, whereas control C2C12 can be clearly distinguished from cells depleted of ERK3 or MK5 at DM2 (Fig. 5A). In addition to confirming the quality of the data, this analysis revealed that loss of ERK3 or MK5 has a clear impact on the transcriptome of differentiating C2C12 myoblasts. We identified a total of 91 and 144 differentially expressed genes (p adjusted < 0.05; log_2_foldchange > 1) in ERK3-depleted and MK5-depleted C2C12 cells, respectively, as compared to control C2C12 in GM (Fig. 5B and Tables S1 and S2). On the other hand, 1,950 and 978 genes were found to be differentially expressed at DM2 (Fig. 5B and Tables S3 and S4). Strikingly, a large proportion of these genes overlapped between ERK3- and MK5-depleted conditions, and only a few genes showed a differential expression when directly comparing the transcriptomes of C2C12 cells depleted of ERK3 or MK5 (Fig. 5B). Variations in gene expression observed upon depletion of ERK3 or MK5 compared to control C2C12 cells were highly correlated, with Pearson coefficients of 0.55 at GM and 0.73 at DM2 (Fig. 5C). These data reinforce the epistatic relationship of ERK3 and MK5 in myogenic differentiation.

**Figure 5.**
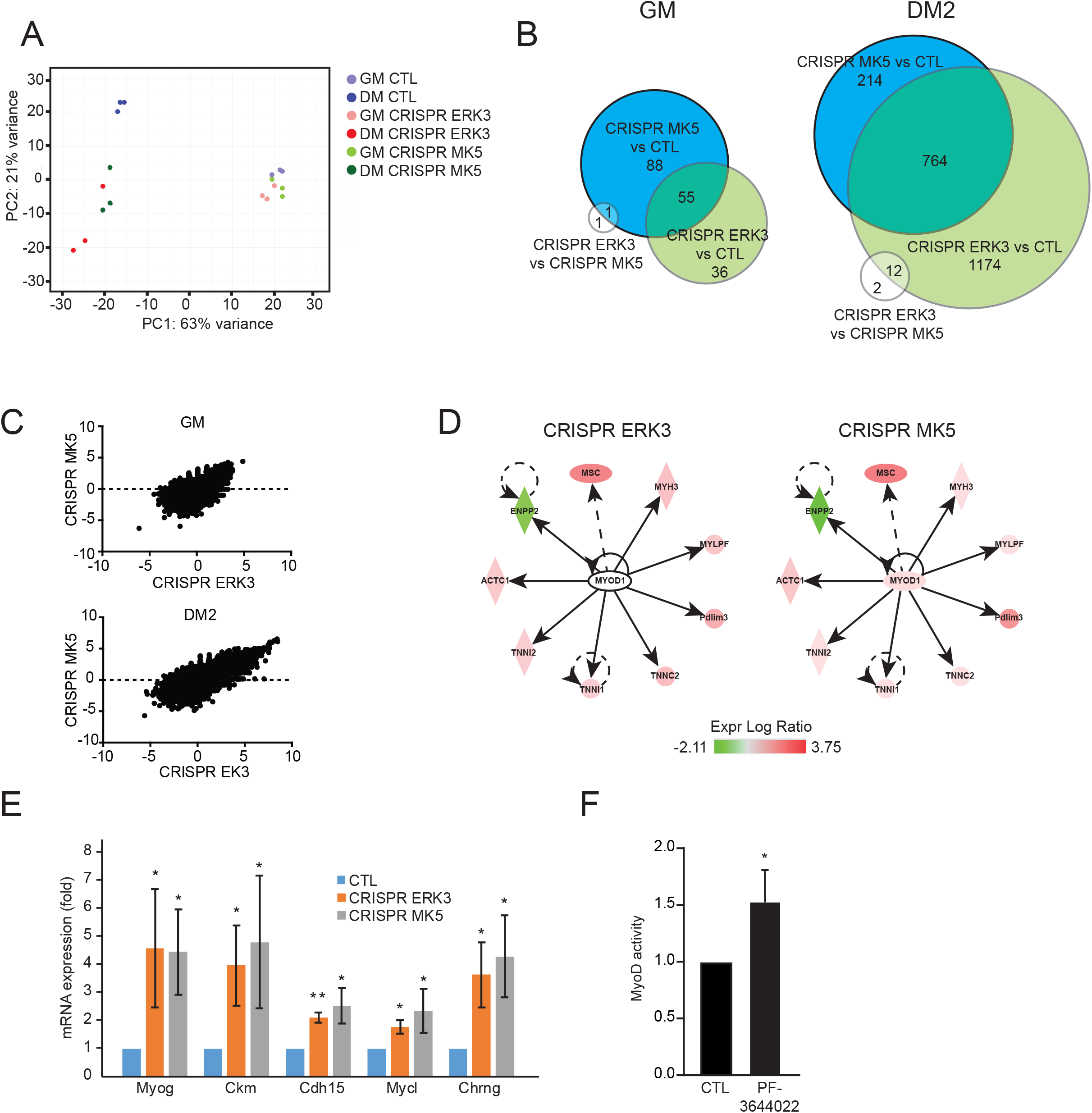
ERK3-MK5 signaling activates MyoD in skeletal myoblasts. (A-D) The transcriptome of control or C2C12 cells depleted of ERK3 or MK5 by CRISPR/Cas9 gene editing was analyzed by RNA-seq at GM and DM2. (A) PCA of the top 5,000 most variable genes. (B) Venn diagrams displaying the number of differentially expressed genes (*p* adjusted < 0.05; log_2_foldchange > 1) in the different experimental conditions. (C) Plot of differentially expressed genes in ERK3-depleted vs MK5-depleted C2C12 cells at GM and DM2. The Pearson coefficient correlation (R value) and *p* value are indicated. (D) Graphical representation of MyoD activation predicted by IPA in ERK3-depleted (*p* = 4.17 × 10^−8^) and MK5-depleted (*p* = 0.031) C2C12 myoblasts in GM. (E) Expression of MyoD target genes was measured by qPCR in control, ERK3-depleted and MK5-depleted C2C12 myoblasts in GM. Values are mean ± SD (n=3). \**p* < 0.05. (F) MyoD transcriptional activity was assayed using a myogenin-luciferase reporter. The cells were treated with 300 nM PF-3644022 for 24 h. Values are mean ± SD (n = 3). \**p* < 0.05.

To identify downstream targets of ERK3 and MK5 signaling, we performed Ingenuity Pathway Analysis (IPA) on the datasets of differentially expressed genes. Upstream regulator analysis identified 55 and 157 transcriptional regulators predicted to contribute to the variations in gene expression observed upon depletion of ERK3 or MK5 in GM (Tables S5 and S6). Interestingly, the myogenic regulatory factor MyoD was predicted to be activated by the loss of ERK3 (*p* = 4.2×10^−8^) or MK5 (*p* = 0.031) in GM (Fig. 5D). We confirmed by qPCR analysis that expression of MyoD target genes is up-regulated in C2C12 myoblasts depleted of ERK3 or MK5 (Fig. 5E).

To directly evaluate the impact of ERK3-MK5 signaling on MyoD transcriptional activity, we used a synthetic reporter containing the MyoD DNA binding site. Pharmacological inhibition of MK5 with PF-3644022 significantly increased MyoD activity in C2C12 myoblasts (Fig. 5F). These findings identify MyoD as an effector of ERK3-MK5 signaling in skeletal muscle differentiation.

### Phosphorylation of FoxO3 by MK5 promotes its degradation and reduces the formation of FoxO3-MyoD complexes

It has been recently reported that MK5 phosphorylates members of the forkhead family of transcription factors, leading to their transcriptional activation (10, 29, 30). FoxO3 was shown to interact with MyoD to orchestrate super-enhancer activity and drive myogenin gene expression during myoblast differentiation (31). We therefore hypothesized that ERK3-MK5 may signal through FoxO3 to modulate MyoD activity and myogenic differentiation. We first confirmed by in vitro phosphorylation assays that recombinant active MK5 directly phosphorylates mouse FoxO3 on Ser214 (Supplementary Fig. S3). To determine if ERK3 or MK5 can physically interact with FoxO3, 293T cells were co-transfected with ERK3, MK5 and FoxO3 and analyzed by co-immunoprecipitation. We found that FoxO3 co-immunoprecipitated with ERK3 when co-expressed with MK5, but not when expressed only with ERK3 (Fig. 6A). When MK5 was immunoprecipitated under the same conditions, FoxO3 co-immunoprecipitated with MK5 regardless whether ERK3 was co-expressed or not (Fig. 6B). These results indicate that ERK3 interacts with FoxO3 through MK5.

**Figure 6.**
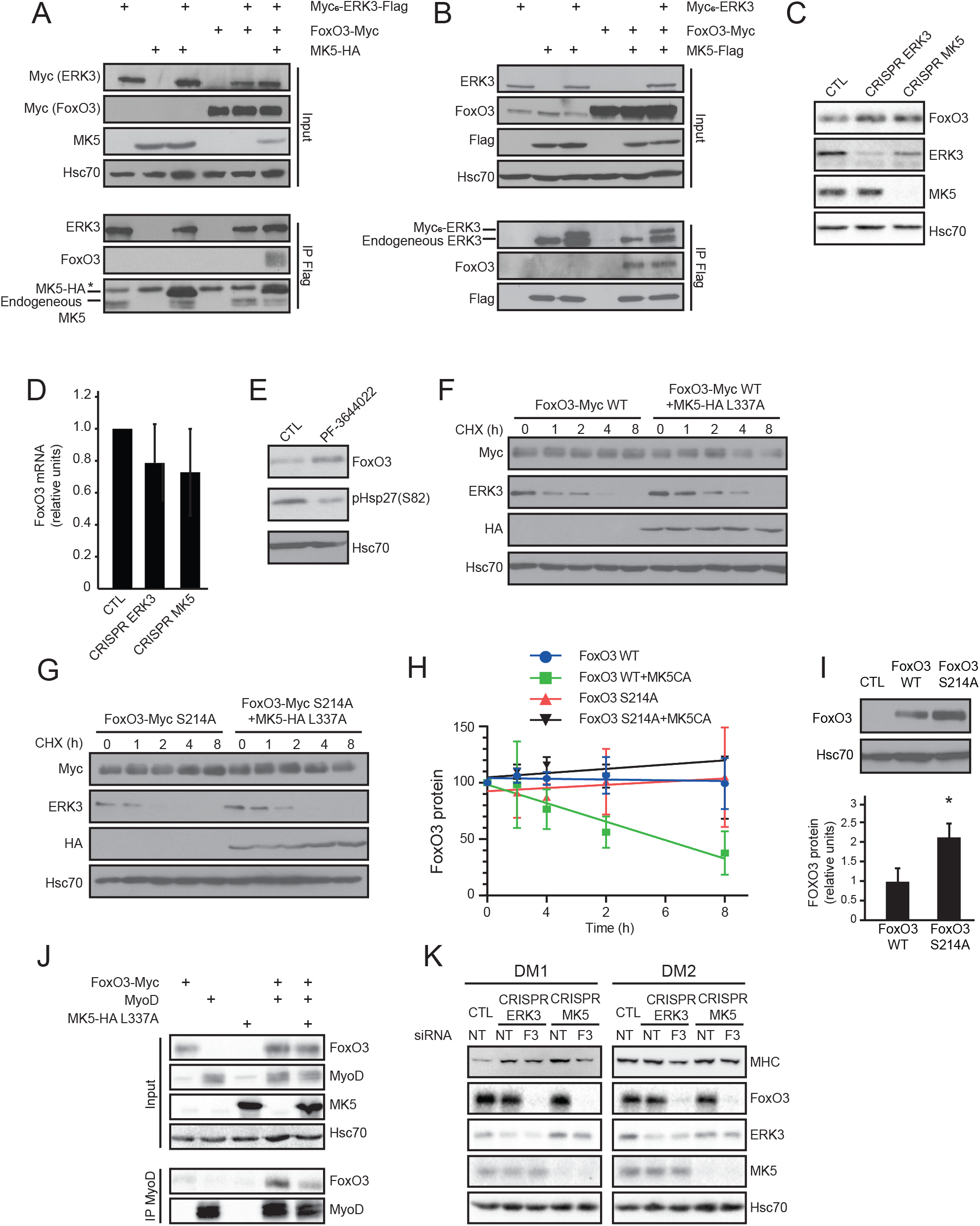
MK5 directly phosphorylates FoxO3 to promote its degradation in skeletal myoblasts. (A and B) Co-immunoprecipitation assays of ERK3 and MK5 with FoxO3. 293T cells were co-transfected with the indicated ERK3, MK5 and FoxO3 constructs and ERK3 (A) or MK5 (B) was immunoprecipitated with anti-Flag antibody. Proteins were analyzed by immunoblotting. (C and D) Expression of FoxO3 protein (C) and mRNA (D) in C2C12 myoblasts after ERK3 or MK5 depletion by CRISPR/Cas9 gene editing. (E) Immunoblot analysis of FoxO3 expression in C2C12 cells treated with 300 nM PF-3644022 for 24 h. Phosphorylation of the MK5 substrate Hsp27 was monitored by anti-phospho-Hsp27(Ser82) immunoblotting to control for MK5 inhibition. (F-H) Cycloheximide (CHX)-chase analysis of FoxO3 stability. 293T cells were transiently transfected with FoxO3 WT or S214A and activated MK5 L337A constructs. After 48 h, CHX was added at 50 μg/ml and the cells were harvested at the indicated times. Proteins were analyzed by immunoblotting. (F and G) Representative immunoblots. (H) Quantification of immunoblotting data (n = 3). (I) C2C12 cells were transiently transfected with FoxO3 WT or S214A mutant and cell lysates were analyzed by immunoblotting. Bottom, quantification of immunoblotting data (n = 3). \**p* < 0.05. (J) Co-immunoprecipitation assay of FoxO3 with MyoD. 293T cells were co-transfected with the indicated FoxO3, MK5 L337A and MyoD constructs and MyoD was immunoprecipitated with anti-MyoD antibody. Proteins were analyzed by immunoblotting. (K) C2C12 myoblasts were depleted of ERK3 or MK5 by CRISPR/Cas9 gene editing. Stable populations were then transfected with FoxO3 siRNAs. After 24 h, the cells were switched to DM to initiate myogenic differentiation. Expression of MHC, FoxO3, ERK3 and MK5 was analyzed by immunoblotting at the indicated times.

Phosphorylation of FoxO3 on Ser215 (human protein, equivalent to Ser214 in mouse) has been proposed to promote its nuclear translocation (10, 30). However, we did not observe any difference in the cellular distribution of WT FoxO3 or FoxO3 S214A mutant in C2C12 myoblasts expressing activated MK5 L337A (Supplementary Fig. S4). We then asked if ERK3-MK5 signaling modulates the levels of FoxO3 in C2C12 myoblasts. Depletion of ERK3 or MK5 by CRISPR/Cas9 editing clearly increased the steady-state protein levels of FoxO3 in C2C12 cells (Fig. 6C) without significant changes in the mRNA levels (Fig. 6D). Similarly, inhibition of MK5 activity with PF-3644022 also increased FoxO3 protein levels in C2C12 cells (Fig. 6E). To determine if MK5 activation affects the stability of FoxO3, we measured the half-life of FoxO3 by cycloheximide-chase experiments. Overexpression of activated MK5 L337A markedly decreased the stability of FoxO3 in 293T cells (Fig. 6 F and H). In contrast, expression of MK5 L337A had no significant effect on the degradation rate of FoxO3 S214A mutant, indicating that Ser214 phosphorylation controls the stability of FoxO3 (Fig. 6 G and H). Accordingly, the steady-state expression of FoxO3 S214A protein was 2-fold higher than WT FoxO3 after transient transfection in C2C12 cells (Fig. 6I).

In agreement with Peng et al. (31), we showed that FoxO3 physically associates with MyoD (Fig. J). Notably, co-expression of MK5 L337A decreased the amount of FoxO3 co-precipitating with MyoD, consistent with the stabilization of FoxO3 protein. To address the functional contribution of FoxO3 to ERK3-MK5 signaling, we silenced the expression of FoxO3 with validated siRNAs in C2C12 myoblasts depleted of ERK3 or MK5 by CRISPR/Cas9 editing. Knockdown of FoxO3 partially rescued the premature differentiation of C2C12 myoblasts induced by depletion of ERK3 or MK5 (Fig. 6K). These results provide evidence that FoxO3 plays a key role in relaying ERK3-MK5 signal during skeletal muscle differentiation.

## DISCUSSION

The physiological roles of the atypical MAP kinase ERK3 remain to be fully investigated. Recently, we reported that ERK3-deficient mice or mice expressing a catalytically-inactive allele of ERK3 show a reduced growth rate postnatally (6). A similar growth retardation phenotype was described in another mouse model of ERK3 deficiency (5). However, the molecular basis of this phenotype was unknown. Body mass increases rapidly during the first three weeks of post-natal life in the mouse, with approximately 50% of the added mass contributed by skeletal muscle (20, 32). In this study, we show that genetic inactivation of ERK3 kinase activity impairs post-natal skeletal muscle growth and adult muscle regeneration, explaining the growth retardation phenotype of these mice. Notably, loss of MK5 leads to similar growth retardation and skeletal muscle development defects. The following observations provide compelling evidence that ERK3 and its substrate MK5 act in a linear signaling pathway to control post-natal skeletal muscle development. First, loss of MK5 phenocopies the muscle growth and regeneration defects of mice expressing ERK3 KD, consistent with a kinase-substrate relationship. Second, genetic depletion of ERK3 or MK5 leads to premature differentiation of skeletal myoblasts in culture. Ectopic expression of the constitutively-active MK5 L337G mutant rescues the differentiation phenotype of ERK3-depleted myoblasts. Third, transcriptomic analyses revealed that the genes that are differentially expressed in ERK3-depleted or MK5-depleted myoblasts compared to control cells are highly correlated.

ERK3-deficient and *Mapk6*^*KD/KD*^ mutant mice have a comparable body weight to WT mice at birth (6). Here, we show that the number of Pax7^+^ satellite cells and the total number of myofibers per TA muscle is similar in adult WT and *Mapk6*^*KD/KD*^ mice. This suggests that specification of Pax7^+^ progenitors and skeletal myoblasts during fetal myogenesis is normal in *Mapk6*^*KD/KD*^ mice, consistent with the idea that ERK3 is dispensable for embryonic and fetal myogenesis. Myofiber number is set around birth in different muscles and does not increase during post-natal life (20). Post-natal muscle growth occurs mainly by individual fiber hypertrophy and is supported by a rapid increase in the number of new myonuclei during the first 3 weeks. The first two weeks are the most active period with the number of myonuclei doubling twice between P3 and P4. Myonuclear accretion ceases by P21 (20). To understand the role of ERK3 in post-natal muscle growth, we analyzed the proliferation/differentiation of skeletal myoblasts. Genetic depletion of ERK3 by CRISPR/Cas9 edition or RNAi induced the premature differentiation of C2C12 myoblasts, associated with a transient increase in myogenin expression. *Mapk6*^*KD/KD*^-derived primary myoblasts also differentiated prematurely compared to WT myoblasts and proliferated at a much slower rate. Genetic or pharmacological inhibition of MK5 similarly induced the precocious differentiation of C2C12 myoblasts, whereas activation of MK5 inhibited differentiation. Importantly, gene expression and cellular pathway analyses predicted that loss of ERK3 or MK5 activates MyoD, which was confirmed by qPCR measurements of target genes and MyoD reporter assay. MyoD-deficient primary myoblasts show defective myogenic differentiation and enhanced proliferation in culture (33), opposite to the phenotype of ERK3-inactive myoblasts. Collectively, our findings suggest the following model for the role of ERK3 during skeletal muscle development (Fig. 7). ERK3 activation of MK5 restrains the activity of MyoD, thereby preventing the premature differentiation of proliferating myoblasts and allowing the accumulation of a pool of myonuclei to sustain myofiber hypertrophy.

**Figure 7.**
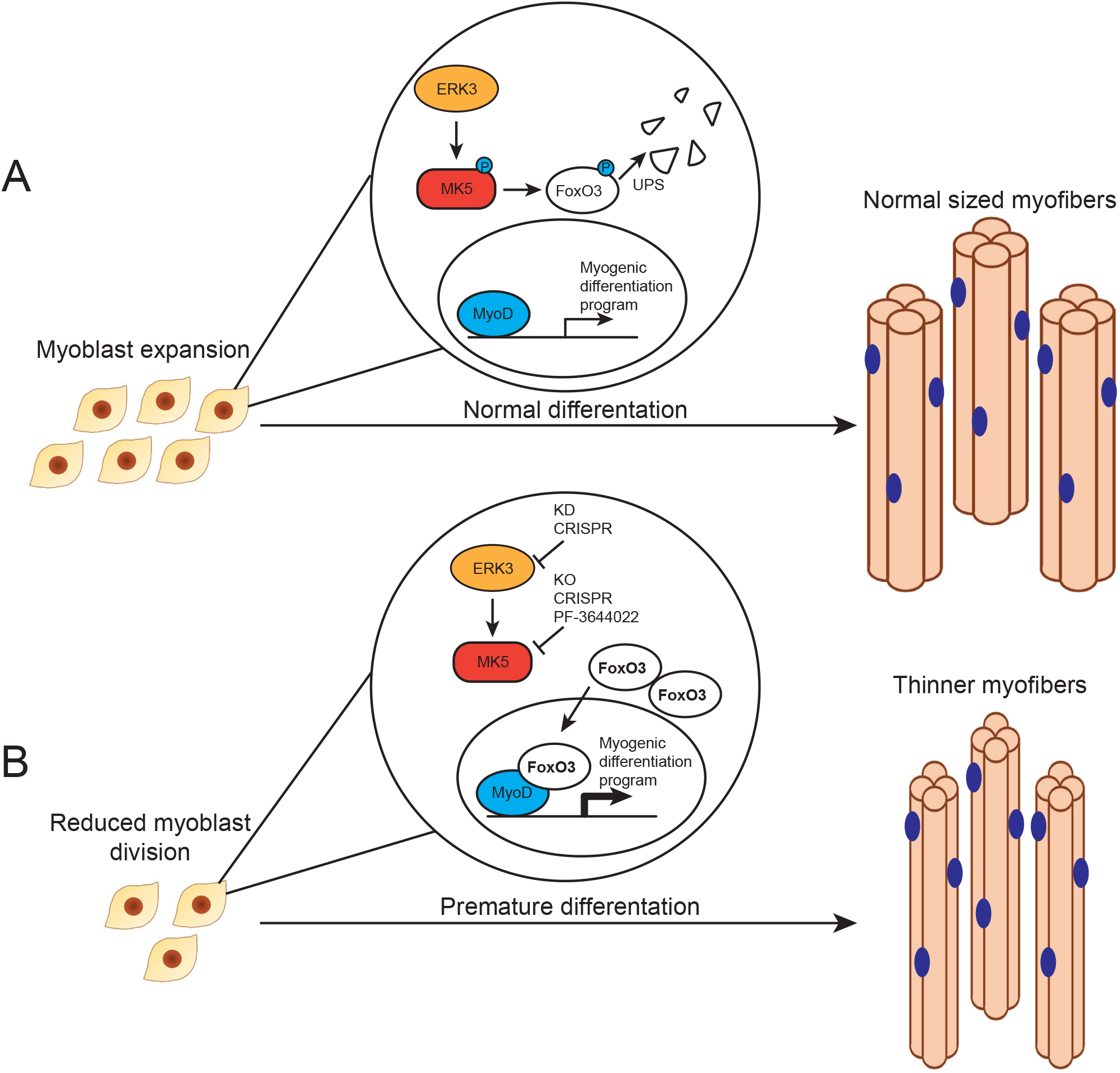
Schematic diagram of the proposed role of ERK3-MK5 signaling in regulating post-natal skeletal muscle development. (A) Normal post-natal myogenic differentiation. (B) Inactivation of ERK3 or MK5 induces premature differentiation of proliferating myoblasts resulting in smaller myofibers. See text for details. KD, kinase-dead; KO, knockout.

This model also explains the defective adult skeletal muscle regeneration observed in *Mapk6*^*KD/KD*^ and *Mapkakpk5^−/−^* mice. Muscle regeneration is dependent on a pool of resident Pax7^+^ satellite cells which, upon muscle injury, re-enter the cell cycle and generate proliferating myoblasts that expand and differentiate to form new myofibers or repair damaged fibers (15, 16). We have shown that the number of Pax7^+^ satellite cells is similar in the TA muscle of WT and *Mapk6*^*KD/KD*^ mice. Decreased proliferation and premature differentiation of myoblasts following muscle injury would reduce the pool of myonuclei available for fusion, resulting in smaller myofibers revealed by a reduction in CSA of centrally nucleated fibers.

A number of MK5 substrates have been identified in recent years, although few have been rigorously validated in vivo (34–36). Among these are members of the FoxO family of transcription factors (37, 38). MK5 phosphorylates human FoxO3 on Ser215 (30) and stimulates the transcriptional activity of FoxO1 and FoxO3 in various cell types (10, 29, 30). Interestingly, a recent study showed that FoxO3 directly binds to MyoD to coordinately orchestrate hot spot assembly and activation of *Myog* super enhancer during C2C12 myoblast differentiation (31). RNAi-mediated depletion of FoxO3 inhibits differentiation of C2C12 myoblasts into myotubes and *Foxo3^−/−^* mice show impaired muscle regeneration after CTX injury (39). These observations prompted us to explore the role of FoxO3 as a downstream effector of ERK3-MK5 signaling in myogenic differentiation. We first confirmed that MK5 directly phosphorylates mouse FoxO3 on Ser214. We further showed that FoxO3 forms a physical complex with ERK3 and MK5 through its interaction with MK5. Of note, FoxO3 was previously identified as a protein interaction partner of ERK3 in a large-scale yeast two-hybrid interaction screen (40). We found that FoxO3 binds to MyoD in C2C12 myoblasts and that activation of MK5 decreases the formation of FoxO3/MyoD complexes. Functionally, RNAi silencing of FoxO3 rescues in part the precocious differentiation of C2C12 myoblasts observed upon inactivation of ERK3 or MK5. These results identify FoxO3 as a direct effector of ERK3-MK5 signaling in the control of skeletal muscle differentiation.

FoxO factors are tightly regulated by a multitude of post-translational modifications that affect protein stability, protein-protein interactions, subcellular localization and transcriptional activity (38, 41). It has been reported that MK5 phosphorylation on Ser215 enhances nuclear localization and DNA binding of FoxO1 or FoxO3 factors (10, 30). However, we failed to observe any impact of Ser215 phosphorylation by MK5 on the nuclear accumulation of FoxO3 in C2C12 myoblasts, suggesting that this regulatory mechanism may be cell type-specific. Instead, we discovered that MK5-mediated Ser214 (mouse numbering) phosphorylation promotes the degradation of FoxO3. Prior studies have shown that ERK1/2 MAP kinases phosphorylate FoxO3 on Ser294, Ser344 and Ser425, increasing its association with Mdm2 and promoting its degradation (42). Oncogenic RAS-dependent up-regulation of casein kinase 1α down-regulates FoxO3 and FoxO4 protein abundance via phosphorylation of Ser318/Ser321 residues in FoxO3 (43, 44). Our findings highlight a new layer to the complex regulation of FoxO factors stability and activity. In a broader perspective, our study adds further support to the importance of MK5 signaling in modulating FoxO activity in different physiological and pathological contexts.

## MATERIAL AND METHODS

### Reagents, plasmids and antibodies

CTX, cycloheximide and PF-3644022 were purchased from Sigma-Aldrich. The plasmids pcDNA3-Myc_6_-ERK3-Flag, pcDNA3-Myc_6_-ERK3, pcDNA3-MK5-HA, pcDNA3-MK5-Flag, pcDNA3-MK5-HA L337A have been previously described (4, 28, 45). pcDNA3-FoxO3-Myc was a kind gift of A. Brunet (Stanford University). pcDNA3-FoxO3-Myc S214A was generated by PCR mutagenesis. To construct the bacterial expression plasmid pGEX-mFoxO3, mouse FoxO3 was amplified by PCR using pcDNA3-FoxO3-Myc as template and subcloned into the EcoRI site of pGEX-KG. All mutations and PCR products were verified by DNA sequencing.

Commercial antibodies were obtained from the following suppliers and used at the indicated concentrations: anti-ERK3 (1/1000; cat. no. ab53277) and anti-myogenin (1/1000; cat. no. 232558) from Abcam; anti-MK5 (1/500; cat. no. sc-46667), anti-Myc (1/1000; cat. no. sc-40) and anti-Hsc70 (1/2000; cat. no. sc-7298) from Santa-Cruz Biotechnology; anti-MHC (1/1000; cat. no. MF20) and anti-Pax7 (1/100) from DSHB; anti-FoxO3 (1/1000; cat. no. 2497) and anti-phospho-Hsp27(Ser82) (1/1000; cat. no.2406) from Cell Signaling Technology; anti-HA (1/1000; cat. no. 901501) from Biolegend; anti-Flag (1/1000; cat. no. F3165), anti-laminin (1/500; cat. no. L9393) and anti-actin (1/1000; cat. no. A4700) from Sigma-Aldrich.

### Mice

*Mapk6*^*KD/KD*^ and *Mapkakpk5^−/−^* mutant mice have been described (6, 46). Mice were housed under specific pathogen-free conditions at the Institute for Research in Immunology and Cancer. For breeding, heterozygous males and females in mixed C57BL/6J x 129/Sv background were intercrossed and littermate controls were used in all studies. All experiments involving animals were approved by the Université de Montréal Institutional Animal Care Committee in compliance with guidelines from the Canadian Council on Animal Care (CCAC).

### Injury-induced skeletal muscle regeneration model

Male mice were used at 10 weeks of age. Mice were injected subcutaneously with 10 mg/kg meloxicam 2 h before CTX injections. The animals were anesthetized with 2% isoflurane, and both TA muscles were injected with 50 µl of 10 µM CTX dissolved in 0.9% saline. TA muscles were harvested 1, 3, 7, 12 and 15 days after injury to monitor the kinetics and extent of muscle regeneration.

### Muscle histopathology analysis

TA muscles were fixed in 10% formalin or frozen in isopentane chilled on a liquid nitrogen bath. H/E sections. Paraffin-embedded muscles were cut in sections of 4 μm and stained with hematoxylin and eosin (H&E). Frozen muscles were cut into 4 µm-thick sections with a cryostat and analyzed by immunofluorescence staining. Briefly, tissue sections were fixed with 10% formalin and permeabilized with cold methanol. After washing with NH_4_Cl, antigen retrieval was performed by incubation with 0.01 M citric acid. Tissues were blocked with M.O.M. reagent (Vector Laboratories), and then incubated with Pax7 and laminin antibodies overnight at 4°C. Bound primary antibodies were detected by incubation with fluorescent secondary antibodies for 1 h at room temperature. Tissue sections were finally stained with DAPI to detect nuclei. Fluorescent images were captured with an Axio Imager Z1 microscope (Zeiss). H&E stained slides were cover-slipped and scanned using the NanoZoomer 2.0-HT digital pathology system (Hamamatsu Photonics). The CSA of myofibers was analyzed with NDP.View2 software (Hamamatsu Photonics).

### Cell culture, transfections and RNA interference

293T cells were cultured in DMEM containing 10% fetal bovine serum (FBS) and antibiotics. C2C12 myoblasts were cultured in DMEM supplemented with 10% FBS and antibiotics (GM). Myogenic differentiation of C2C12 cells was induced by replacing GM with DM (DMEM with 2% horse serum) when the cells reached 90-95% confluence (4). Satellite cell-derived primary myoblasts were cultured in Ham’s F10 media containing 20% FBS, 1% penicillin-streptomycin, and 2.5 ng/mL basic fibroblast growth factor (Wisent). Myogenic differentiation was induced by replacing the medium with 50% Ham’s F10, 50% DMEM low-glucose, 1% penicillin-streptomycin, and 5% horse serum.

293T cells were transiently transfected with polyethylenimine and 12 µg of plasmid DNA per 100-mm culture dish (3:2 ratio). The cells were harvested 36 h post-transfection. C2C12 cells were transfected using Lipofectamine 2000 reagent according to the manufacturer’s specifications (Invitrogen). For siRNA transfections, C2C12 cells were transfected with RNAiMAX (Invitrogen) according to the manufacturer’s instructions. For shRNA lentiviral infections, 293T cells were transfected with pMD2/VSVG, pMDLg/pREE, pRSV/Rev, and the indicated pLKO.1 construct. After 48 h, virus-containing culture media was filtered and used to infect C2C12 cells. Polyclonal populations of infected cells were obtained by selection with puromycine. Lentiviral shRNA clones targeting *Mapk6* and *Makpakp5* were from the TRC1 shRNA library (Sigma-Aldrich): TRCN0000023199 for *Mapk6* and TRCN0000024197 for *Makpakp5. FoxO3* SMARTpool ON-TARGETplus siRNAs (L-040728-00-0005) was purchased from Dharmacon.

### CRISPR/Cas9 gene edition

For CRISPR/Cas9 gene editing, C2C12 cells plated in 3.5-cm petri dish were transfected with 2μg of PX459 vector (47) expressing sgRNA sequences targeting *Mapk6* or *Mapkapk5* gene. The following sgRNA sequences were used: *Mapk6* sg1, 5’-CACCGCCATTGGGCTGCGGAGGCAA-3’ (forward) and 5’-AAACTTGCCTCCGCAGCCCAATGG-3’ (reverse); *Mapk6* sg2, 5’-CACCGCTTTACTGGAAGAGCATGCC-3’ (forward) and 5’-AAACGGCATGCTCTTCCAGTAAAG-3’ (reverse); *Mapkapk5* sg1, 5’-CCACGTAGAAGAATATAGTATCAAT-3’ (forward) and 5’-AAACATTGATACTATATTCTTCTAC-3’ (reverse); *Mapkapk5* sg2, 5’-CCACGCAGTGCAAACCGTTCTTGAG-3’ (forward) and 5’-AAACCTCAAGAACGGTTTGCACTGC-3’ (reverse). Stable populations were generated by selection in 4 μg/ml puromycin for 2 days and used thereafter.

### Isolation of primary myoblasts

Primary myoblasts were isolated as previously described (48). Hind limb muscles of wild type and *Mapk6*^*KD/KD*^ mice were dissected, and the tissues were dissociated using the gentleMACS Dissociator (Miltenyi Biotec). Briefly, muscles were minced in 5 ml of collagenase/dispase solution (2.5 U/ml), and placed into gentleMACS C tubes where they were attached to heating elements and incubated with a custom digestion program: 3 min at 60 rpm (clockwise) for further dissociation of tissue chunks, 9 min at 30 rpm (counterclockwise) for digestion, 10 cycles of 5 s at 360 rpm (clockwise and counterclockwise alternatively) for trituration, and 12 min at 30 rpm (counterclockwise) for secondary digestion. The suspension of mononuclear cells was filtered through a 50-µm nylon filter, washed with FACS buffer (5% FBS and 1 mM EDTA in PBS), and stained with phycoerythrin(PE)-conjugated anti-Sca-1 (clone D7, cat. no. 553108, BD Biosciences), anti-CD45 (clone 30-F11, cat. no. 12-0451-83, BD Biosciences), anti-CD31 (clone 390, cat. no. 12-0311-81, BD Biosciences), anti-CD11b (clone M1/70, cat. no. 12-0112-81, BD Biosciences), APC-conjugated anti-Itga7 (clone R2F2, cat. no. 67-0010-10, AbLab), and 7-AAD (Biolegend). Satellite cells were isolated by FACS using a gating strategy gated based on forward scatter (FSC) and side scatter (SSC) profiles, cell viability (7-AAD negative cells), negative lineage selection in PE for removal of Sca-1-, CD45-, CD31- and CD11b-positive cells, and positive lineage selection in APC for Itga7. A small fraction of prospectively isolated cells was cytospun onto slides immediately after isolation with Cytospin 4 (Thermo Scientific) at 500 rpm for 10 min, and analyzed for purity and expression of myogenic markers by immunofluorescence staining. The remaining cells were suspended in growth medium and plated on collagen-coated petri dishes. All FACS experiments were performed on a BD FACSAria Fusion.

### Immunofluorescence analysis of myoblast differentiation

C2C12 cells were plated on glass coverslips coated with 2% gelatin and myogenic differentiation was induced by switching to DM. Cells were washed twice with ice-cold PBS, and then fixed with 3.7% paraformaldehyde in PBS for 20 min at room temperature. After quenching in 0.1 M glycine in PBS for 10 min, the cells were permeabilized by incubation in 0.1% Triton X-100 in PBS for 2 min at room temperature, and then blocked in 1% bovine serum albumin in PBS for 30 min at 37 °C. For detection of MHC, the cells were stained for 1 h at room temperature with anti-MHC primary antibody, followed by incubation with Alexafluor 488 anti-mouse secondary antibody for 1 h. Samples were washed and counterstained with DAPI to detect nuclei. A similar procedure was used for immunofluorescence analysis of satellite cell-derived primary myoblasts, except that the blocking step was performed for 60 min with a 5% goat serum, 2% bovine serum albumin solution, and primary antibody incubation was performed overnight at 4°C.

### Cell proliferation assays

C2C12 cell proliferation was measured using the WST1 colorimetric assay (Roche). Briefly, C2C12 myoblasts were seeded into 96-well plates in triplicate and cultured for the indicated times. Cell proliferation was assayed by reading the absorbance at 450 nm with a reference filter at 620 nm. The proliferation of satellite cell-derived primary myoblasts was monitored using an IncuCyte S3 live-cell imaging system (Essen Bioscience). Briefly, satellite cell-derived primary myoblasts were plated in 96-well collagen-coated plates (5,000 cells/well) and, after 12 h, the plates were placed in the Incucyte for 4 days. The system automatically took pictures every 6 h and performed unbiased analysis of cell count and cell confluence using the IncuCyte cell-by-cell analysis software module.

### Quantitative PCR analysis

Real-time quantitative PCR was performed as previously described (6).

### Gene expression profiling

To identify downstream targets of ERK3-MK5 signaling, we analyzed the transcriptomes of C2C12 cells depleted of ERK3 or MK5 by CRISPR/Cas9 gene editing at GM and at DM2. Total RNA was extracted from cell lysates using RNAeasy purification kit (Qiagen). The quality of RNA was assessed on the Agilent 2100 Bioanalyzer and the RNA was quantified by QuBit fluorometry. Libraries (500 ng total RNA) were prepared using the KAPA Hyperprep mRNA-Seq Kit (KAPA Biosystems). cDNA fragments were ligated to indexed library adapters (Illumina) prior to PCR amplification (11 cycles). Purified libraries were normalized by qPCR using the KAPA Library Quantification Kit (KAPA Biosystems) and diluted to a final concentration of 10 nM. Sequencing was performed on the Illumina Nextseq 500 using the Nextseq 500 High Output Kit (75 cycles) with 2 pM of the pooled library (read length 1 x 80 bp). Around 24-31 M single-end pass-filter reads was generated per sample. Library preparation and sequencing was made at the Institute for Research in Immunology and Cancer Genomics core facility.

Alignment of RNA-seq reads was performed with the STAR aligner and differential gene expression was calculated with DESeq2 using the RNA Express software suite (Illumina). A *p* value < 0.05 was used as threshold for significant changes in gene expression. Cellular pathway analysis was performed with Ingenuity Pathway Analysis (IPA) software (Qiagen).

### MyoD reporter assay

MyoD transcriptional activity was assayed using a myogenin-luciferase reporter (49). C2C12 myoblasts were seeded in 12-well culture plates at 3×10^4^ cells per well. The day after, the cells were transfected with myogenin-luciferase (1 µg) and Renilla luciferase (30 ng) plasmids using lipofectamine 2000 (ThermoFisher). After 6 h, the medium was replaced with fresh GM supplemented or not with PF-3644002. Firefly and Renilla luciferase activities were measured 24 h after transfection.

### Immunoblot analysis and immunoprecipitation

Cell lysis, immunoblot analysis, and immunoprecipitation were performed as described previously (4). For immunoprecipitation experiments, 500 µg of lysate proteins were incubated with 2 µl of anti-Flag monoclonal antibody pre-adsorbed on protein A-Sepharose beads (cat. no. A2220, Sigma-Aldrich) overnight at 4°C.

### In vitro phosphorylation assays and mass spectrometry analysis

Recombinant wild type or mutant mouse FoxO3 proteins were produced in *E. coli* BL-21 strain and purified on glutathione-agarose beads. For in vitro phosphorylation assays, 50 ng of purified recombinant active MK5 (SignalChem) was incubated with 15 µg of purified recombinant FoxO3 in 50 µl of phosphorylation buffer (40 mM Tris-HCl (pH 7.5), 20 mM MgCl_2_, 1 mg/ml bovine serum albumin, 1 mM dithithreitol and 40 µM ATP) at 37 °C for 30 min. Samples were reconstituted in 50 mM ammonium bicarbonate with 10 mM TCEP (ThermoFisher Scientific), and vortexed for 1 h at 37°C. Chloroacetamide (Sigma-Aldrich) was added for alkylation to a final concentration of 55 mM. Samples were vortexed for another hour at 37°C. One microgram of trypsin was added, and digestion was performed for 8 h at 37°C. Samples were dried down and solubilized in 5% acetonitrile-0.2% formic acid. The samples were loaded on a 1.5 ml pre-column (Optimize Technologies). Peptides were separated on a custom-made reversed-phase column (150-µm i.d. by 200 mm) with a 56-min gradient from 10 to 30% acetonitrile-0.2% formic acid and a 600-nl/min flow rate on an Easy nLC-1000 connected to a Q-Exactive HF (ThermoFisher Scientific). Each full mass spectrometry (MS) spectrum acquired at a resolution of 60,000 was followed by tandem-MS (MS-MS) spectra acquisition on the 15 most abundant multiply charged precursor ions. Tandem-MS experiments were performed using collision-induced dissociation at a collision energy of 27%. The data were processed using PEAKS X (Bioinformatics Solutions) and a Uniprot mouse database (16,977 entries). Mass tolerances on precursor and fragment ions were 10 ppm and 0.01 Da, respectively. Fixed modification was carbamidomethyl (C). Variable selected posttranslational modifications were oxidation (M), deamidation (NQ), phosphorylation (STY). The data were visualized with Scaffold 4.3.0 (protein threshold 99%, with at least 2 peptides identified and a false-discovery rate (FDR) of 1% for peptides).

## ACKNOWLEDGMENTS

We thank Kim Lévesque and Mélania Gombos for expert assistance with animal experimentation, Patrick Gendron for help with bioinformatics data analysis, Raphaelle Lambert for qPCR analysis, Julie Hinsinger for histology assistance, and Éric Bonneil for MS analysis. This work was supported in part by grants from the Canadian Institutes for Health Research and the Cancer Research Society to S. Meloche. S. Meloche held the Canada Research Chair in Cellular Signaling.

