## Supplementary Figures for "ERK3-MK5 signaling regulates myogenic differentiation and muscle regeneration by promoting FoxO3 degradation"

Figure S1

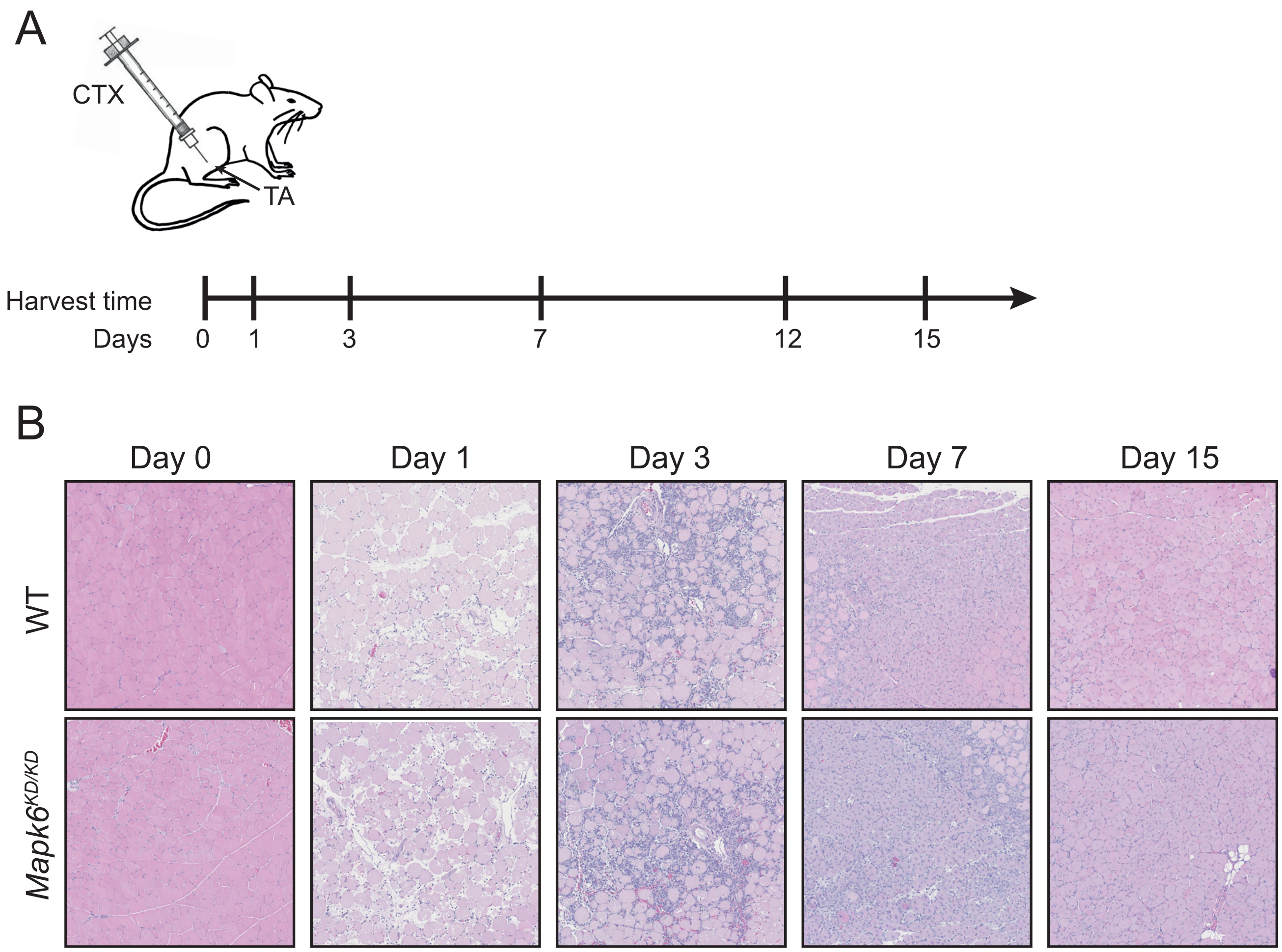


Figure S1. Model of cardiotoxin (CTX)-induced muscle injury and regeneration. (A) Schematic diagram of the experimental model of CTX-induced acute muscle injury. CTX is injected in the tibialis anterior (TA) muscle of 10-week old mice. The TA muscles are harvested at the indicated times and subjected to histopathological analysis. (B) Representative micrographs of H&E stained TA sections at different times after CTX injection showing regeneration of the injured muscle after 15 days.

#### Figure S2

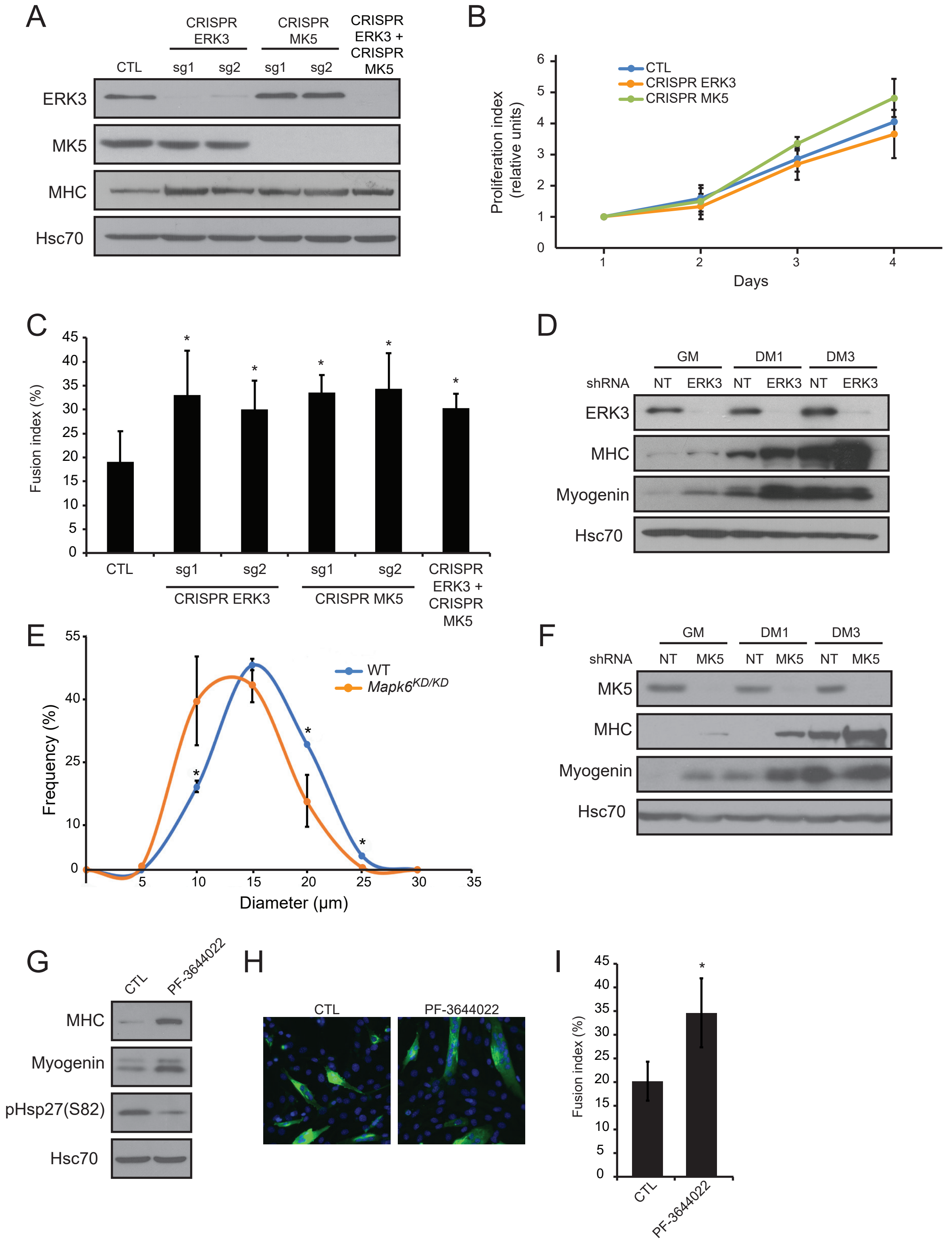


Figure S2. Genetic or pharmacological inactivation of ERK3-MK5 signaling induces premature differentiation of skeletal myoblasts. (A-C) Analysis of C2C12 myoblasts depleted of ERK3 and/or MK5 by CRISPR/Cas9 gene editing. C2C12 cells were transfected with empty vector (CTL) or PX459 vector expressing sgRNA sequences (sg) targeting Mapk6 and/or Mapkapk5 gene. Stable populations were generated by puromycin selection and used thereafter. The cells were grown in GM medium and switched to DM medium to initiate myogenic differentiation. (A) Expression of MHC, ERK3 and MK5 was analyzed by immunoblotting at DM2. (B) Cell proliferation was measured using the WST-1 assay. Values are expressed as fold increase and represent the mean +/- SD of 4 replicates. (C) Quantification of fusion index of differentiating C2C12 myoblasts at DM3. Values are mean +/- SD ($n \geq 3$). *$p < 0.05$. (D) C2C12 myoblasts were infected with non-target (NT) shRNA or ERK3 shRNA-expressing lentiviruses. Expression of MHC, myogenin and ERK3 was analyzed by immunoblotting at the indicated times. (E) Satellite cell-derived primary myoblasts were isolated from WT and Mapk6KD/KD mice. Quantitative analysis of myotube diameter after 4 days of differentiation. Values are mean +/- SD (n = 3). *$p < 0.05$. (F) C2C12 myoblasts were infected with non-target (NT) shRNA or MK5 shRNA-expressing lentiviruses. Expression of MHC, myogenin and MK5 was analyzed by immunoblotting at the indicated times. (G-I) Effect of the MK5 inhibitor PF-3644022 on C2C12 myoblast differentiation. C2C12 cells were treated with 300 nM PF-3644022 for 24 h in GM prior to switching to DM in the continuous presence of inhibitor. (G) Expression of MHC and myogenin was analyzed by immunoblotting at DM2. Phosphorylation of the MK5 substrate Hsp27 was monitored by anti-phospho-Hsp27(Ser82) immunoblotting to control for MK5 inhibition. (H) Immunofluorescence staining of MHC expression at DM3. (I) Quantification of fusion index of differentiating C2C12 myoblasts at DM3. Values are mean +/- SD (n =4). *$p < 0.05$.

Figure S3

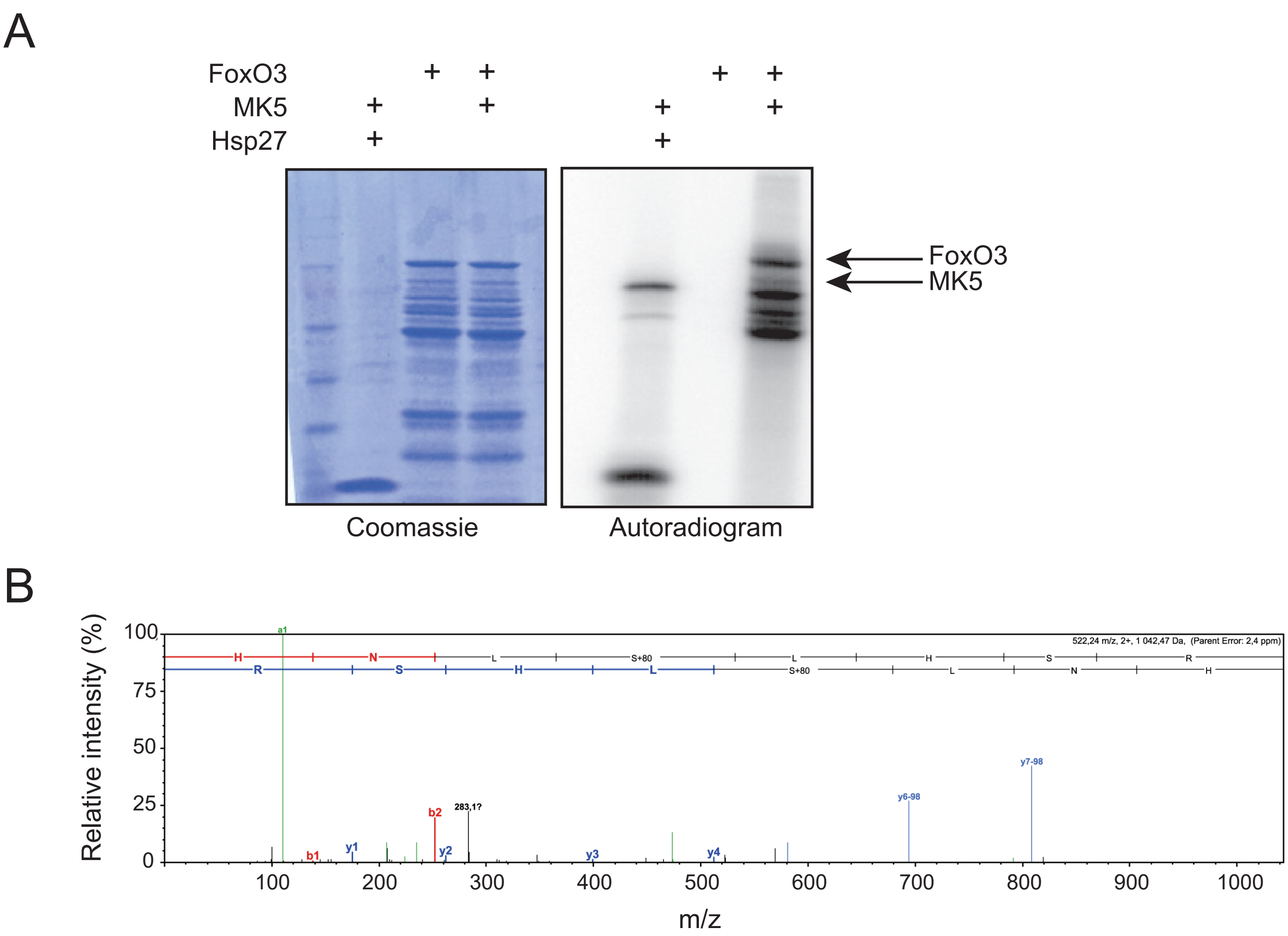


Figure S3. MK5 directly phosphorylates mouse FoxO3 on Ser214. (A) In vitro phosphorylation assay. GST-FoxO3 was produced in E. coli and incubated with recombinant active MK5 in kinase assay buffer supplemented with 40 µM ATP and 1 µCi [γ-32P] ATP for 30 min at 30oC. Phosphorylation was analyzed by autoradiography. (B) MS/MS spectrum showing the phosphorylation of mouse FoxO3 on Ser214.

### Figure S4

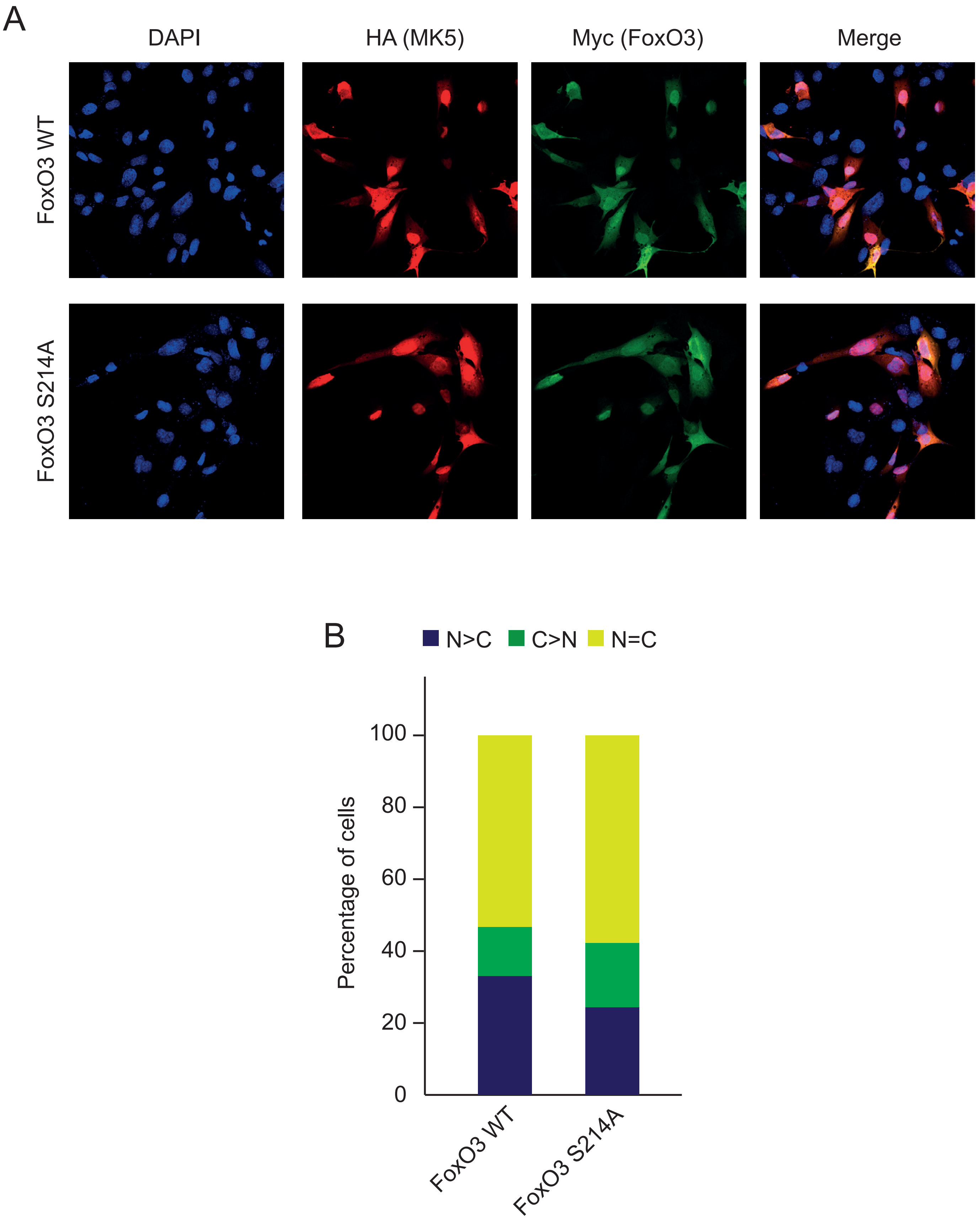


Figure S4. Phosphorylation of human FoxO3 on Ser214 does not affect its subcellular localization in C2C12 myoblasts. (A) C2C12 cells were co-transfected with MK5 L337A and either WT or S214A mutant FoxO3. Localization of MK5 L337A and FoxO3 was analyzed by immunofluorescence after 24 h. (B) Quantification of FoxO3 cellular localization. Between 50 and 100 cells were counted in each experiment. Values are mean of 4 independent experiments. N, nuclear, C, cytoplasmic.
